# multiSMD – a Python toolset for multidirectional Steered Molecular Dynamics

**DOI:** 10.1101/2025.07.21.665906

**Authors:** Katarzyna Walczewska-Szewc, Beata Niklas, Wiesław Nowak

## Abstract

Understanding the direction-dependence of molecular interactions is critical for elucidating biological processes such as protein-protein binding, ligand dissociation, and mechanotransduction. While steered molecular dynamics (SMD) simulations enable the study of force-induced transitions, conventional single-direction approaches may overlook anisotropic responses inherent to biomolecular systems. Here, we present multiSMD, a Python-based tool that automates the setup and analysis of multi-directional SMD simulations in NAMD and GROMACS. By systematically probing forces along multiple spatial vectors, multiSMD captures direction-dependent phenomena—such as varying energy barriers or structural resilience—that remain hidden in traditional SMD. We demonstrate the utility of our approach through three distinct applications: (i) anisotropic unbinding in a protein-protein interaction, (ii) ligand dissociation pathways dependent on pulling direction, and (iii) force-induced remodeling of intrinsically disordered regions. multiSMD streamlines the exploration of mechanical anisotropy in biomolecules, offering a computational framework to guide experiments (e.g., AFM or optical tweezers) and uncover mechanistic insights inaccessible to single-axis methods.

Availability and implementation: multiSMD is freely available at https://github.com/kszewc/multiSMD

## 1. Introduction

Forces and their temporal dynamics are fundamental to the functionality and regulation of living systems, even at the molecular scale. Proteins experience and exert forces that drive essential biological processes, including folding, ligand binding, transport, and enzymatic catalysis. These forces arise from molecular interactions—electrostatic, van der Waals, hydrogen bonding, and hydrophobic effects—and their modulation over time enables responsiveness to environmental stimuli and biochemical signals. Critically, many of these processes are inherently anisotropic, meaning that their mechanical and dynamic properties vary depending on spatial direction due to the complex, non-spherical structures of biomolecules. Understanding the time-dependent and direction-dependent nature of these forces is crucial for elucidating the mechanisms such as mechanotransduction, allosteric regulation, and binding/unbinding events.

The development of advanced methodologies to investigate molecular forces has significantly enhanced our understanding of their role in biological systems. Atomic force microscopy (AFM), particularly in conjunction with force spectroscopy^1^, has enabled the direct measurement of piconewton-scale forces in single molecules, revealing insights into ligand binding^2^, antibody-antigen interactions^3^, protein unfolding^4^, or enzymatic catalysis^5^. This high-resolution imaging technique bases on scanning the sample surface with a very sharp tip and measuring the cantilever deflection thus giving information on the tip-sample interaction. Similarly, optical and magnetic tweezers apply controlled forces to molecules (often via attached beads) enabling precise manipulation and study of DNA mechanics, motor protein motility, or receptor-ligand dissociation^6,7^. While single-molecule force spectroscopy experiments performed with AFM or tweezers are immeasurably useful, they are also very challenging. Problems with proper immobilization of biomolecules during sample preparation, single-molecule identification, and discrimination of non-specific interactions make this technique unsuitable for high-throughput investigations^8^.

Computational methods, particularly molecular dynamics (MD) simulations provide a complementary approach, serving as a “computational microscope” for molecular biology^9^. The time evolution of molecular systems at atomic resolution is predicted in molecular dynamics (MD) simulation, where forces acting on each atom of investigated biomacromolecule are calculated to update their position in space. In steered molecular dynamics (SMD), time-dependent external forces are applied to the system along the preselected coordinate to accelerate transitions between energy minima in the free energy landscape. This enables studying biophysical processes that require time scales not accessible for classical MD, such as protein (un)folding^10^, transport across a membrane^11^, identification of ligand binding pathways^12,13^, or elucidation of the dynamics of big protein complexes^14,15^. SMD also serves as a tool for comparing binding affinities of small-molecule ligands to their target proteins by measuring the force required to pull them out of their binding sites^16^, thus providing valuable insights into the inhibitory potential of drug candidates. While traditional SMD simulations focus on a single pulling direction, many biological processes—such as the anisotropic response of proteins to mechanical stress or the directional dependence of ligand unbinding pathways—are best studied through multi-directional force probing^17^. Such an approach can reveal subtle differences in energy landscapes, hidden transition states, and direction-dependent resistance to deformation, which might otherwise remain undetected in conventional single-axis simulations. Multi-directional SMD simulations can thus complement and guide experiments like AFM and optical/magnetic tweezers by exploring force-dependent processes from multiple angles before committing to labor-intensive experimental setups.

To bridge this gap, we developed multiSMD, a tool that automates the preparation of inputs for multi-directional SMD simulations. This Python-based script generates a series of batch files enabling SMD simulations in popular MD engines (NAMD and GROMACS) based on the provided protein complex structure (see Figure 1a). To facilitate data postprocessing, multiSMD comes with a dedicated script *Analysis*.*py* for analyzing the obtained results. This script allows the user to extract and visualize data relevant to the simulated systems. By enabling systematic exploration of force responses along multiple spatial vectors, multiSMD facilitates a more comprehensive understanding of anisotropic biological processes such as protein mechanostability, direction-dependent unbinding mechanisms, or the mechanical roles of structural motifs in large complexes.

**Figure 1.**
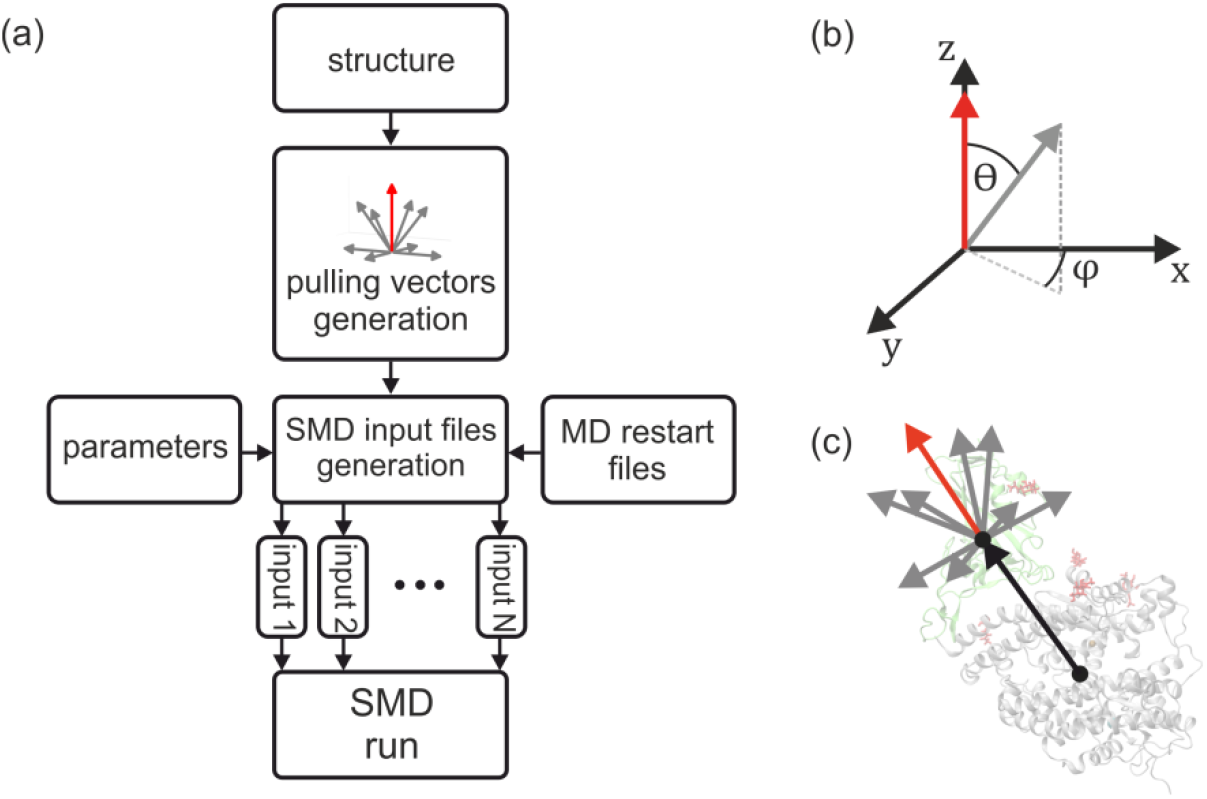
(a) Flowchart illustrating the operation of multiSMD. (b) Angles describing the successive generated pulling vectors. (c) The directions of external force application in parallel SMD simulations of the test system of the S-protein-ACE.

## 2. Implementation

The multiSMD program is written in Python 3 and maintained on GitHub (kszewc/multiSMD). The flowchart of our program is shown in Figure 1a. Provided the Cartesian coordinates of protein complex atoms in the PDB format, the program computes the principal axis of “pulling,” which serves as a vector connecting the centers of mass of the fixed and pulled proteins (see Figure 1c,b). Drawing from this principal axis, multiSMD generates a comprehensive set of vectors, characterized by variations in theta and phi angles within spherical coordinates. Each vector within this set denotes a unique direction for stretching the molecular system during SMD simulations. Notably, users retain the flexibility to adjust the sampling density of this force vector space, with default settings encompassing three angles (0, 45, and 90 deg) in the theta coordinate and four angles (0, 90, 180, 270 deg) in the phi coordinate. This configuration yields a total of nine distinct pulling directions, effectively covering a hemisphere. Upon execution, multiSMD generates an output directory containing the input files and subdirectories corresponding to the “pull” directions. Each subdirectory contains an appropriately prepared input files for NAMD/gromacs and a bash script to run the given SMD simulation.

The additional scripts (Analysis.py for NAMD and Analysis_gro.py for gromacs) allows for extracting essential SMD data from MD output files and obtained trajectories. This includes information about the change in pulling force over time, which allows us to assess the point of the maximum pulling force. Additionally, the script calculates the dependence of pulling force on the distance between centers of masses of two defined atom groups, allowing us to observe the system’s response to the applied force directly. Furthermore, using the MDAnalysis library, one can analyze the obtained trajectories to monitor formation and breakage of hydrogen bonds between the protein fragments of interest. All these quantities are plotted using the Matplotlib graphics library. The Example output from the multiSMD analysis script, showing the force variation over time, force versus distance, and the time-dependent change in hydrogen bond count due to directional pulling is shown as a supplementary Figure S1.

For visualization purposes, the program generates a Tcl script to be used in Visual Molecular Dynamics (VMD) software^18^, which draws a bunch of vectors representing the directions of pulling.

## 3. Results and discussion

We applied our method on three protein systems, each representing distinct application: (i) protein-protein interaction, (ii) protein-ligand interaction, and (iii) interaction involving intrinsically disordered regions.

### 3a. Case study I: Investigating the forces required to disrupt SARS-CoV-2 S protein – ACE2 complex in multiple directions

We first evaluated the efficacy of our multiSMD program by applying it to the COVID-19 relevant protein system. A key stage of viral infection is the interaction between viral spike proteins and human angiotensin-converting enzyme 2 (ACE2) receptor.^19^ Current drug discovery strategies for COVID-19 focus on weakening these specific protein-protein interactions by blocking the ACE2 receptor or the viral Spike protein to prevent infection. We thus proceed to measure the forces required to disrupt this complex considering their anisotropy.

We used the structure of the receptor-binding domain (RBD) of the spike protein bound to ACE2^20^ (PDB ID 6M0J, see Supplementary Figure S2). Using microsecond long MD simulation of the complex^21^ we were able to extract a structurally stable fragment of the complex interface (see Supplementary information for details). We also performed *in silico* mutagenesis of the ACE2 receptor, focusing on three mutations, S19W, T27W, and N330Y (see Figure 2b), previously shown to enhance SARS-CoV-2 S-RBD binding^22^, to investigate their impact on SMD pulling forces. The structures of the SARS-CoV-2 S-RBD bound to the ACE2 mutants reveal that the increased binding affinity is mainly due to van der Waals interactions created by the aromatic side chains of W19, W27, and Y330.

**Figure 2.**
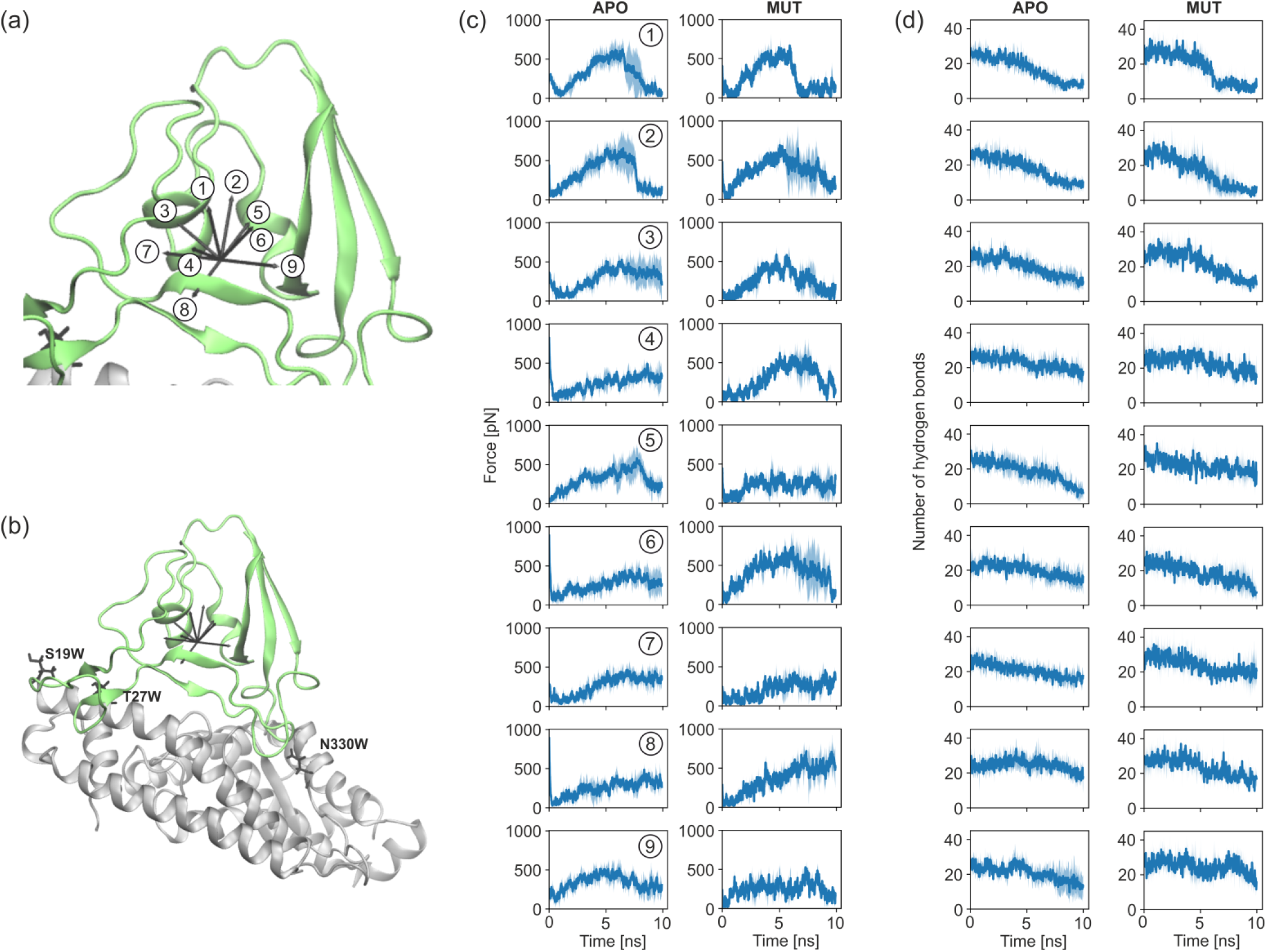

After equilibration and 0.25 ns of classical MD simulation using NAMD performed to stabilize the system, we run our multiSMD.py script to generate inputs for SMD simulations in nine directions of pulling (see Figure 2a). We run 5 replicas of 10 ns SMD in each direction for non-modified complex (APO) and mutated system (MUT).A detailed description of the methods and investigated system is provided in the Supplementary Information.

We based our analysis on a calculation of (i) the changing number of hydrogen bonds when pulling the RBD in various directions (Figure 2d), and (ii) forces required to disrupt the complex (Figure 2c). In a simulation time as short as 10 ns, we observed a significant anisotropy in the complex’s response to external forces. In a system without modifications (APO), pulling in all directions resulted in the reduction of hydrogen bonds. However, when mutations were introduced, the number of hydrogen bonds was constant upon pulling in some directions (see MUT (45, 180), (45, 270), and (90, 90) in Figure 2d). Greater forces were also required to disrupt the mutated complex, although not in all tested directions. These results suggest stronger interactions between RBD and ACE2 when substitutions to residues with aromatic side chains are present, which may enhance virulence. One should bear in mind that the simulations were performed using limited interacting parts of both proteins (not a full system) so the allosteric effects were not possible to capture.

### 3b. Case study II: The comparison of ATP unbinding from Kir6.1 and Kir6.2

Inward-rectifying potassium (Kir6.x) proteins (Figure 3a) are the pore-forming subunits of ATP-sensitive potassium channels (KATP). These channels link membrane potential with intracellular ATP levels^23^. Different combinations of Kir6.1 and Kir6.2 and sulfonylurea receptor (SUR1, SUR2a, and SUR2b) subunits generate various KATP subtypes with distinct tissue distributions and functions^24^. Despite significant sequence and structural similarity, Kir6.1 and Kir6.2 isoforms differ in their sensitivity to ATP^25,26^. The ATP-binding site in both isoforms is highly conserved, with nearly identical residues involved in ligand interaction, as confirmed by available cryo-EM structures (eg. 6C3P^27^ and 7MIT^25^) and our unbiased MD simulations. Snapshots of ATP bound to Kir6.1 and Kir6.2 are shown in Figure 3c,e. Figure S4 illustrates the frequency of close contacts between individual residues and ATP throughout the unbiased simulation.

**Figure 3.**
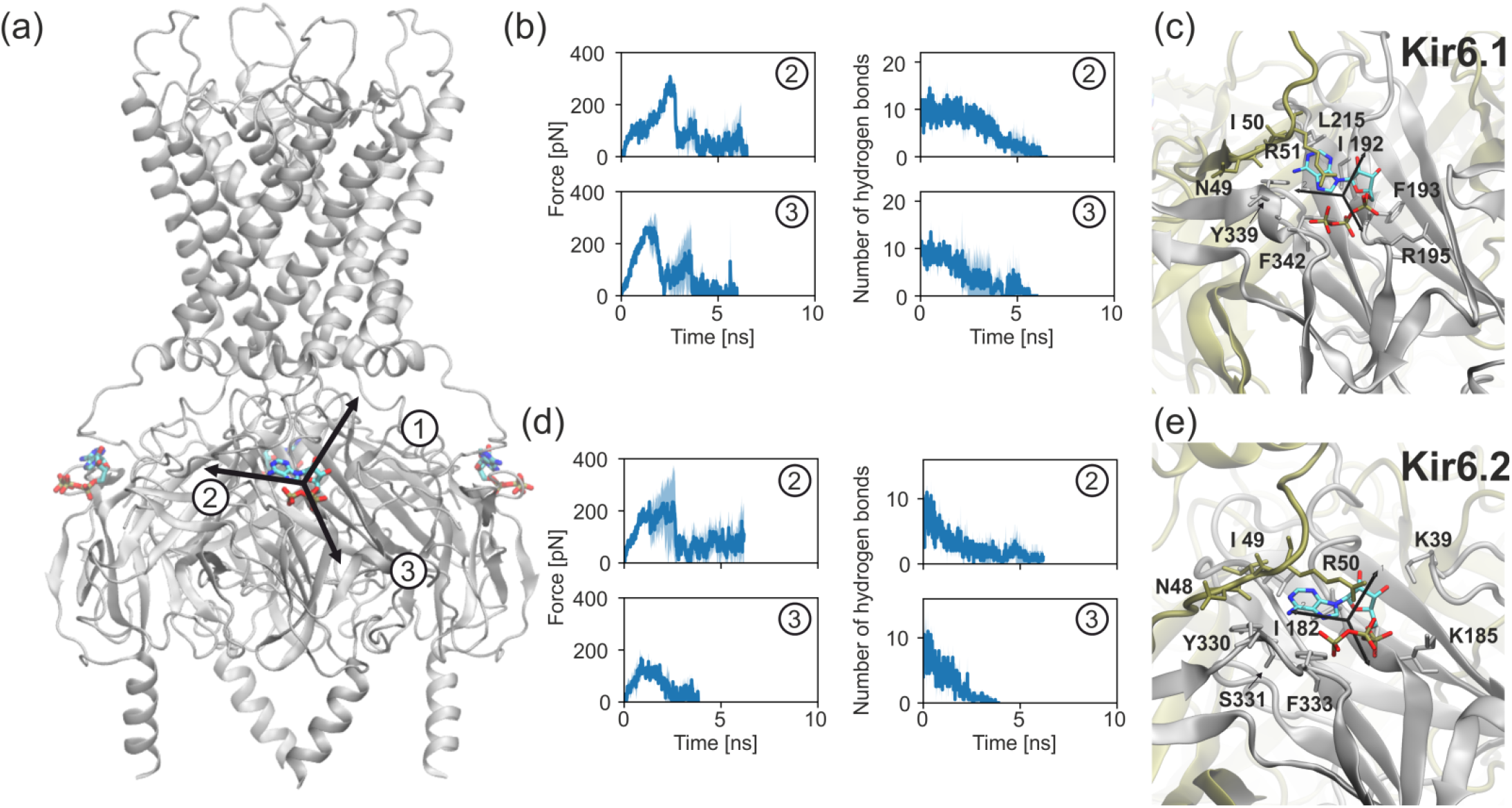
Direction-dependent ATP unbinding from Kir6.1/Kir6.2 channels. (a) Structural overview of Kir6.x tetramer (cartoon) with bound ATP (sticks). (b,d) Force profiles during ATP extraction along direction 2 and direction 3, showing Kir6.1 (b) versus Kir6.2 (d). (c,e) MD snapshots of ATP (cyan) binding sites in Kir6.1 (c) and Kir6.2 (e), highlighting key residues (sticks).

Experimental studies have reported notable differences in ATP-binding between the isoforms despite the high conservation of the ATP-binding site. One distinguishing feature is the substitution of R195 in Kir6.1 with K185 in Kir6.2. Both residues are positively charged and critical for ATP binding, yet the substitution may subtly influence ligand interaction. To explore this further, we applied our multiSMD method to evaluate the forces required to extract ATP from the binding pocket at a constant velocity of 0.0005 nm/ps. Starting from equilibrated Kir6.1 and Kir6.2 systems, we pulled ATP along three directions, recording unbinding forces until complete ligand dissociation.

We excluded the first pulling direction from analysis, as the ATP trajectory led to the cell membrane, which has no physiological rationale. Our results show significant differences in forces required for pulling along the third direction, where Kir6.1 requires 1.5 times greater force than Kir6.2. In turn, pulling along the second direction yields similar force profiles for both isoforms (Figure 3b,d). Notably, direction 3 leads ATP toward the region involving the R195/K185 substitution. Analysis suggests that R195 in Kir6.1 forms stronger electrostatic interactions with the triphosphate moiety of ATP than K185 in Kir6.2, which may explain the observed difference. For the third pulling direction, forces required to pull the ATP molecule out of its binding site were higher for Kir6.1 than for Kir6.2, and a greater amount of time was required to break the hydrogen bonds between the Kir6.1 protein and the ligand.

These findings are consistent with the hypothesis that Kir6.1 may exhibit tighter ATP binding compared to Kir6.2, at least in the context of the isolated Kir6 tetramer. However, it is important to note that ATP sensitivity in the full KATP channel complex (including the SUR subunit) is influenced by additional factors, such as interactions with SUR, the presence of Mg-nucleotides, and PIP2 interactions, which are not captured in our simplified model. For instance, Kir6.1-containing channels are known to require Mg-nucleotides to open and exhibit lower open probability compared to Kir6.2-containing channels, despite potentially stronger ATP binding to Kir6.1^25,26,28^. Thus, while our simulations provide insights into the differences in ATP-binding mechanics between Kir6.1 and Kir6.2, they do not fully explain the physiological differences in ATP sensitivity observed in the complete KATP channel complex.

These preliminary findings highlight the potential of our multidirectional SMD method for identifying subtle yet functionally significant differences in ligand binding and interaction sites. Further energetic analysis, as well as simulations incorporating the full KATP channel complex, will be needed to confirm these results and provide deeper insights into the interplay between ATP binding, channel gating, and regulation by SUR subunits

### 3c. Case study III: KNt Release from SUR2B Pocket in Vascular KATP Channels

The tight interaction between Kir6.x and SUR subunits, which involves both their interfacial contacts and the interactions of disordered regions, is a recently discovered/described feature of KATP channels^15,25,29–31^. The N-terminus of Kir6.2 in pancreatic KATP channels and Kir6.1 (termed KNt) in vascular KATP channels enhances this interaction by inserting into a relatively distant pocket in SUR1 and SUR2B, respectively. This interaction, through a not yet fully understood mechanism, leads to channel closure. In cryo-EM structures of closed KATP channels, densities corresponding to the N-terminus of Kir6 can be observed within this pocket^25,31^. However, the N-terminus is absent from the pocket in open-channel structures where SUR is dimerized, suggesting a physiological insertion and removal process^32^. This indicates that KNt should effortlessly exit the pocket when required.

The process of KNt release from the SUR2B pocket is enigmatic. Its directionality is unclear because KNt, an intrinsically disordered region (IDR), lacks a defined position outside the pocket. This makes it an ideal system for testing our multiSMD approach. A simulation system comprises SUR2B with the KNt region of Kir6.1 inserted into the pocket (Figure 4a), representing a fragment of the vascular KATP channel. We performed unbiased MD simulations to equilibrate the system. Two systems were constructed: one without ligands and another with glibenclamide in the pocket. Glibenclamide stabilizes the KNt position and supports the inward-open conformation of SUR2B, corresponding to the KATP channel’s closed state^31^.

**Figure 4:**
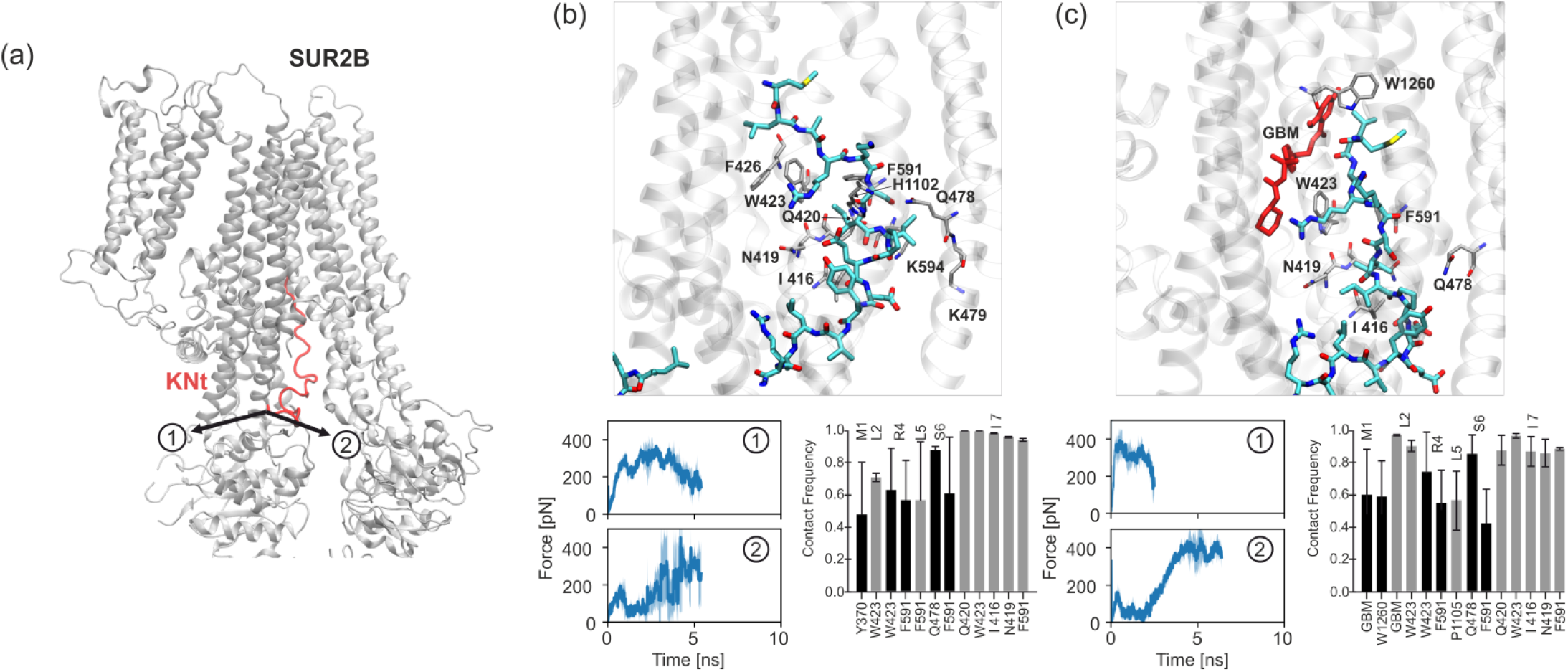
Multi-directional analysis of KNt release from SUR2B. (a) System overview with two tested pulling directions (arrows). (b) Without glibenclamide: contact frequencies (right), binding pocket snapshot (upper), and extraction force profiles (left). (c) With glibenclamide: equivalent analyses showing stabilized KNt interactions. Force curves (blue) demonstrate direction-dependent differences in extraction mechanics.

The frequency of close contacts between KNt and SUR residues in the unbiased simulation is presented as bar plots in Figures 4b and 4c (for systems with and without glibenclamide, respectively). Snapshots illustrating the position of KNt within the pocket and surrounding residues are also shown. We identified two possible pulling directions for KNt for the SMD simulations, denoted as directions 1 and 2 (Figure 4a). Artificial pulling forces were applied to the proximal part of KNt (residues 20-22), allowing us to evaluate which direction requires less force and, consequently, suggests a more straightforward release pathway.

The profiles of forces required to extract the distal KNt region (residues 1-10) from the pocket in a function of simulation time, are shown in blue plots in Figures 4b and 4c for systems without and with glibenclamide, respectively. The graphs cut off at the point where KNt fully exits the SUR1 pocket. Significant differences between the first and the second pulling directions were observed. For direction first, a force of approximately 400 pN was initially required to overcome strong interactions between SUR’s E1196 and KNt’s K24 as well as E1173 and R23. These residues form stable connections/interactions that must be disrupted for the KNt tail to exit the pocket. In this case, the force is applied tangentially to these interactions, which is less effective than applying it perpendicularly. Therefore, pulling along the second direction was initially easier, with resistance increasing as the distal KNt region began exiting the pocket.

Notably, the presence of glibenclamide slightly increased the force required for KNt extraction, particularly in the second direction. While these findings suggest potential pathways and interactions affecting KNt release, more precise methods, such as umbrella sampling, are necessary for a detailed energetic characterization of the process.

## 4. Conclusions

Biological systems exhibit fundamental anisotropy in their mechanical responses, yet most computational tools study molecular interactions along single directions. Our multiSMD approach addresses this limitation by enabling systematic, multi-directional force probing of biomolecular systems. Through three representative case studies, we demonstrated how this method provides unique insights into direction-dependent phenomena.

In the SARS-CoV-2 spike-ACE2 system, we observed that stabilizing mutations increased mechanical resistance preferentially along specific pulling vectors, explaining their enhanced binding affinity. Comparative studies of Kir6.1/Kir6.2 channels revealed isoform-specific ATP binding mechanics, with up to 1.5-fold differences in unbinding forces depending on pulling direction. The analysis of KNt extraction from SUR2B further demonstrated how intrinsically disordered regions exhibit path-dependent release mechanisms modulated by small molecules.

The multiSMD toolkit simplifies these analyses through automated workflow generation for major MD packages (NAMD, GROMACS) and integrated analysis tools. While the current implementation focuses on basic force profiling, the modular design permits future expansion to advanced sampling techniques. As an open-source resource, multiSMD aims to make anisotropic force analysis accessible to both specialists and non-experts.

This approach complements existing experimental techniques by identifying critical pulling directions before laborious AFM or optical tweezer experiments. Looking ahead, we anticipate applications in rational drug design, where understanding direction-dependent binding mechanics could optimize therapeutic compounds. The case studies presented here establish multiSMD as a practical tool for probing the directional complexity inherent to biological macromolecules.

## Data Availability Statement

The multiSMD software is made available under the Apache 2.0 license, ensuring accessibility and adherence to non-commercial terms of use. The program, accompanied by comprehensive documentation, is hosted on GitHub at the following repository: https://github.com/kszewc/multiSMD.

## Supporting Information Available

Supporting Information is available free of charge at https://pubs.acs.org/.

- Example multiSMD analysis outputs showing force profiles and structural changes
- Molecular model of the truncated ACE2-spike protein complex
- Conformational dynamics analysis (RMSF) of full vs. truncated systems
- Residue-specific contact frequencies between Kir6.1/Kir6.2 and ATP

## Acknowledgements

We acknowledge Polish high-performance computing infrastructure PLGrid for awarding this project access to the LUMI supercomputer, owned by the EuroHPC Joint Undertaking, hosted by CSC (Finland) and the LUMI consortium through PLL/2023/04/016512. We thank Maciej Lewandowski for his work on early stages of this project, Viktorija Dujmović for her work during Toruń Students Summer Program in Exact Sciences and Dr Lukasz Peplowski for fruitful discussions. We are also grateful to Prof. Show-Ling Shyng for her expertise and discussions on KATP channels.

The work on multiSMD has been supported by NCU in Torun, Poland within ANTICO/IDUB project (WN).

Authors declare no conflict-of-interest.

