## Supplementary File for "multiSMD – a Python toolset for multidirectional Steered Molecular Dynamics"

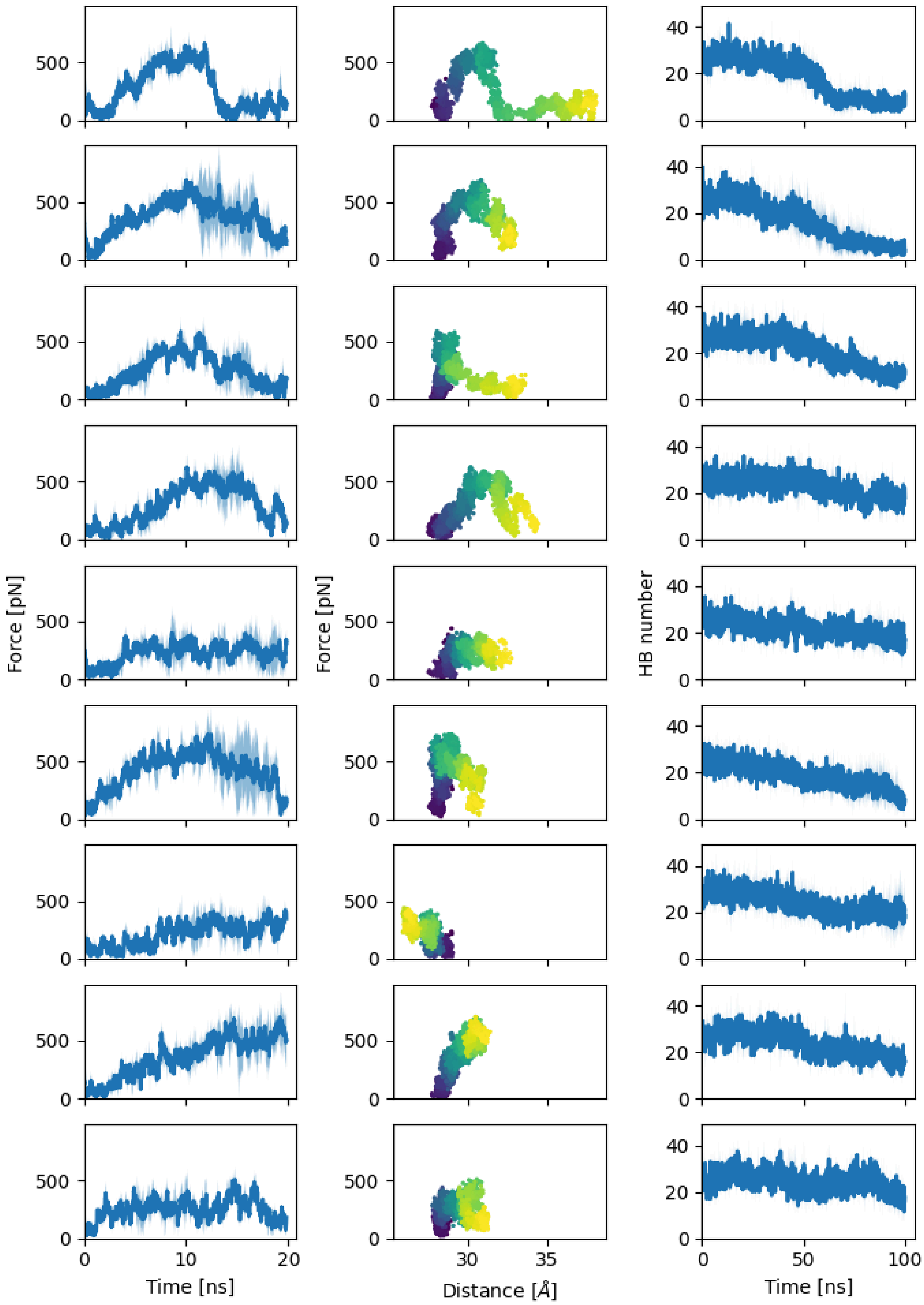

Figure S1: Example output from the multiSMD analysis script, showing the force variation over time (left), force versus distance between two selected anchor points (colored by simulation time progression, middle), and the time-dependent change in hydrogen bond count due to directional pulling (right).

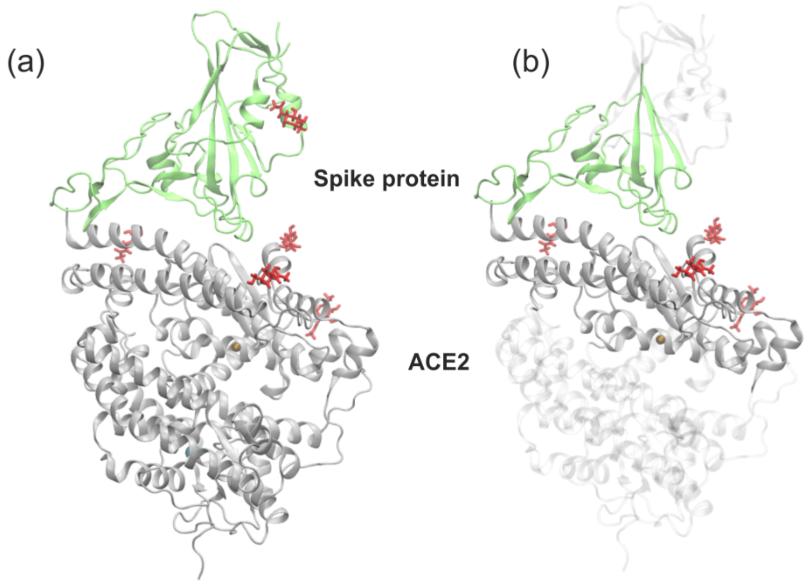


Figure S2: The truncated molecular system of the ACE2-S-protein complex used in our simulations. The ACE2 receptor is shown in gray and the viral spike protein is shown in green. Glycosylation sites are highlighted in red.

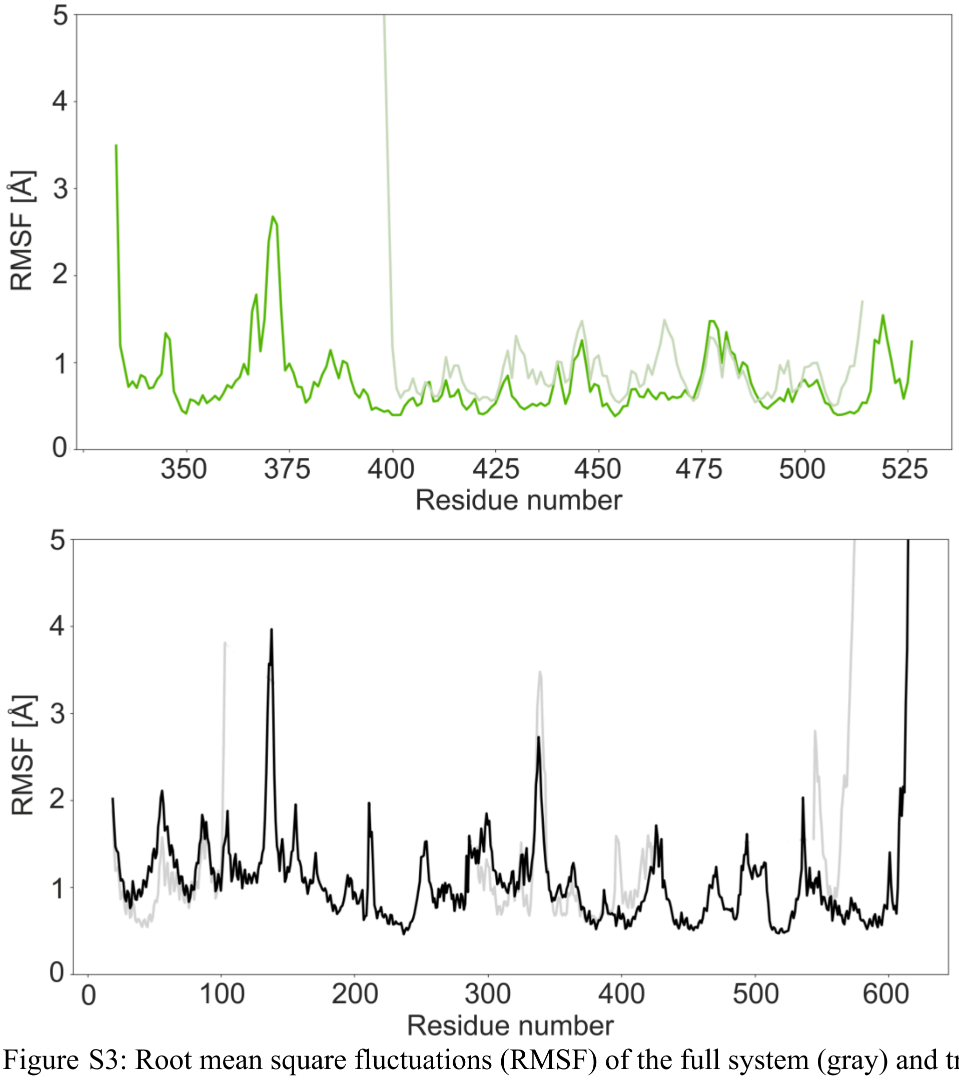


Figure S3: Root mean square fluctuations (RMSF) of the full system (gray) and truncated system (green and black) for the viral S-protein (upper panel) and ACE2 receptor (lower panel).

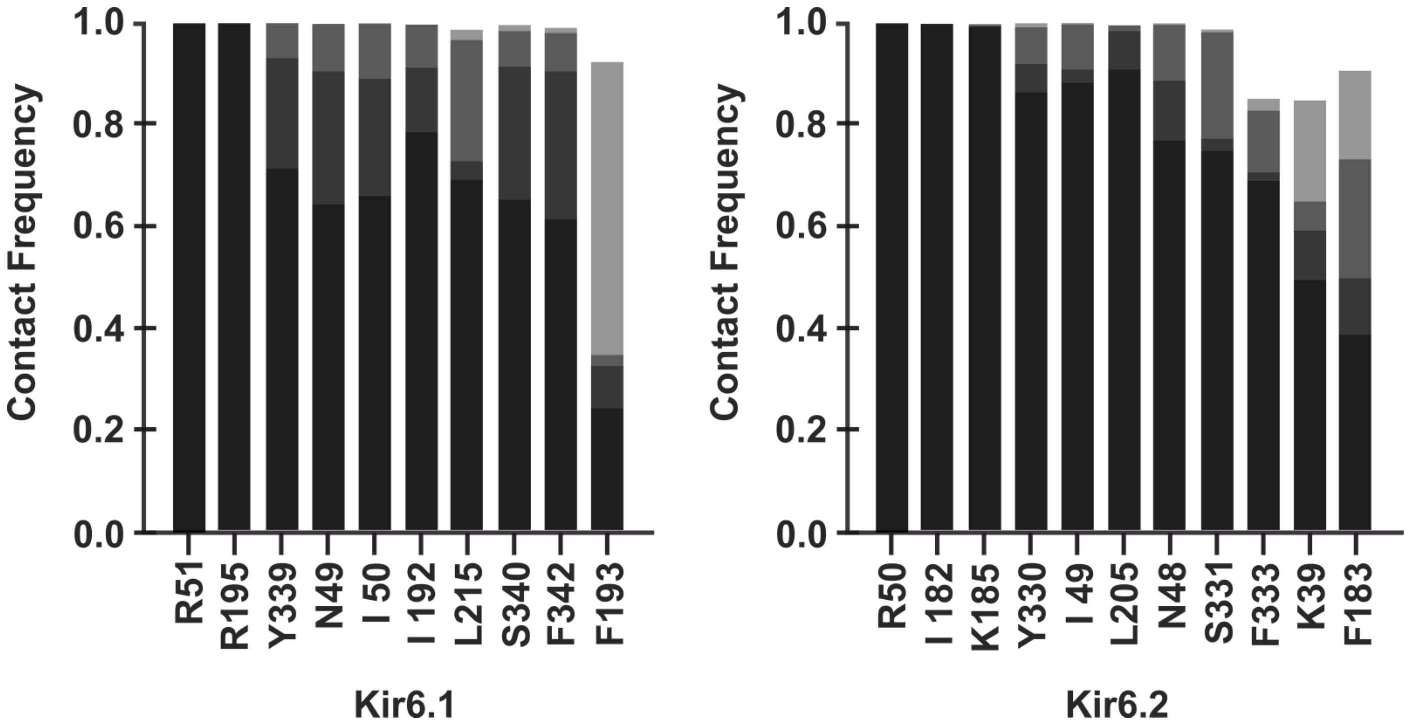


Figure S4: Frequency of close contacts between Kir6.1/Kir6.2 residues and ATP (docked at the inhibitory binding site) during unbiased molecular dynamics simulations.
